# Effects of PTMs on Tau Protein Aggregation: Insights from HCG and Atomistic MD Simulations

**DOI:** 10.64898/2026.05.22.727278

**Authors:** Adelie Louet, Jan Stuke, Lisa Pietrek, Michele Vendruscolo, Gerhard Hummer

## Abstract

Post-translational modifications (PTMs) of the tau protein are increasingly recognized as pivotal regulators in the onset and progression of tauopathies, such as Alzheimer’s disease (AD). To systematically evaluate the structural and functional consequences of specific PTMs, we generated and analyzed seven distinctly modified variants of the tau-K32 construct. These included phosphorylation at Ser202/Thr205, phosphorylation at Ser258/Ser262/Ser356, full phosphorylation at all reported Ser/Thr sites, acetylation at Lys274/Lys281, acetylation at Lys280, full acetylation at all sites, and an unmodified control. Selection of PTM sites was guided by prior experimental literature. By incorporating fully modified tau models, we assessed the global impact of widespread modifications on structural properties and aggregation behavior. Our findings establish a comparative framework for understanding how discrete and cumulative PTMs modulate tau aggregation and provide mechanistic insight into PTM-induced tau dysfunction relevant to neurodegenerative diseases.

## Introduction

Tau is the predominant microtubule-associated protein (MAP) in mature neurons, functioning alongside MAP1 and MAP2^1^. It exists in six isoforms in the adult human brain, all encoded by a single gene on chromosome 17 through alternative splicing. Tau plays a key role in stabilizing microtubules by binding to their surface and promoting tubulin polymerization. Unlike most cytosolic proteins, tau is classified as an intrinsically disordered protein (IDP), characterized by a lack of stable secondary structure and high conformational flexibility^2^.

In neurodegenerative diseases, collectively referred to as tauopathies, tau undergoes pathological alterations, most notably abnormal hyperphosphorylation. This leads to the formation of tau filaments that accumulate into neurofibrillary tangles (NFTs), a hallmark of Alzheimer’s disease. NFTs are composed of paired helical filaments (PHFs) and straight filaments (SFs)^1,3–5^ . Tauopathies also include disorders such as frontotemporal dementia with Parkinsonism linked to chromosome 17 (FTDP-17), Pick disease, corticobasal degeneration, dementia pugilistica, and progressive supranuclear palsy. Hyperphosphorylation is widely believed to be a primary driver of tau dysfunction. Despite being disordered, tau retains its flexibility even when bound to tubulin, forming a “fuzzy” complex that permits multiple binding modes^6^.

Structurally, tau contains key domains that regulate its aggregation propensity, particularly the proline-rich and pseudo-repeat regions. PTMs such as phosphorylation and acetylation are critical modifiers of tau structure and function. Phosphorylation alters tau’s interaction with microtubules and modulates its tendency to aggregate. For instance, phosphorylation at Ser262 has been shown to impair microtubule binding and reduce fibrillization^7^. Experimental work has demonstrated that triple phosphorylation at Ser258, Ser262, and Ser356 abolishes fibril formation in the K18 fragment, suggesting this combination may exert neuroprotective effects. Conversely, phosphorylation at Ser202, Thr205, and Ser208 promotes aggregation^8,9^. Acetylation, on the other hand, disrupts tau’s electrostatic interactions and hydrogen bonding^10^. Acetylation at Lys280 impairs microtubule binding and increases the pool of soluble, aggregation-prone tau. Increased acetylation at Lys274 and Lys281 has been observed in AD patients and is associated with functional impairments in transgenic mouse models^11^.

Molecular Dynamics (MD) simulations have become an important addition to observe the characteristics of structural ensembles of tau. MD simulations captured the structural effects of mutations in the proline-rich regions associated with altered microtubule interaction and aggregation propensities^12^. All-atom simulations of tau fragments have revealed transient secondary structure elements, long-range intramolecular contacts, and conformational preferences that are otherwise difficult to access experimentally due to tau’s disordered nature. These can be translated to identify mechanisms, such as oligomerization^13^, as well as how various post-translational modifications can enhance or attenuate certain secondary structures of the peptide^14^. More specifically, simulations have shown the relationship between certain point mutations can impact the biological mechanism of tau binding to microtubules and the stability of the resulting complex^15^ .

To investigate the structural impact of these PTMs, we generated seven systems for comparative analysis by all-atom MD simulations: tau-K32 variants phosphorylated at Ser202/Thr205, Ser258/Ser262/Ser356, and fully phosphorylated at all known sites **(Figure 1a)**; acetylated tau-K32 variants at Lys274/Lys281, Lys280, and fully acetylated at all potential sites **(Figure 1b)**; along with an unmodified tau-K32 serving as the control (**SI Table 1**). Each post-translational modification (PTM) site was selected based on experimentally validated data from the literature. To further explore the impact of extensive modifications on tau structure and behaviour, we also included fully phosphorylated and fully acetylated chains. These hyper-modifications profoundly alter the charge profile of tau-K32 (**Figure 1**).

**Figure 1:**
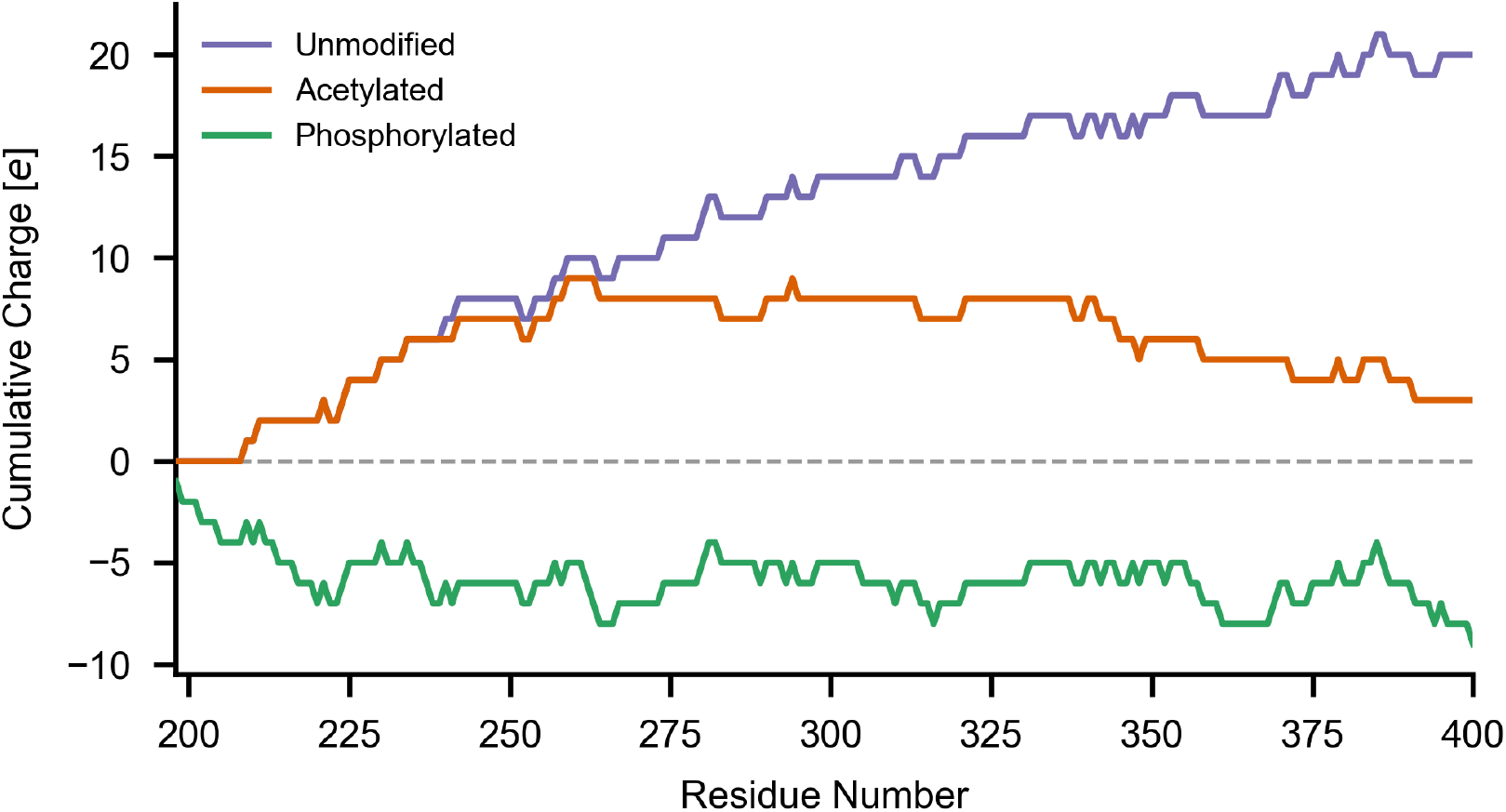
Electrostatic effects accumulated along the tau-K32 peptide chain when fully phosphorylated and acetylated versus unmodified. The cumulative charge obtained by summation along the chain illustrates how electrostatics influence the overall charge distribution from the N-terminus to the C-terminus.

## Results

### Generation of Tau Structures Using Hierarchical Chain Growth (HCG)

Due to tau’s disordered nature, available experimental structural data is limited. To generate representative starting conformations, we employed the Hierarchical Chain Growth (HCG) algorithm^16,17^, which constructs ensembles of IDPs through iterative assembly of overlapping structural fragments. HCG-derived structures served as the basis for our molecular dynamics (MD) simulations. HCG does not consider long-range interactions and, consequently, initial analysis revealed that HCG produces more extended conformations compared to MD simulations (**Figure 2)**. Interestingly, even without considering long-range interactions, HCG showed the same trend as the simulations-more compact tau-K18 chains in response to hyperphosphorylation.

**Figure 2:**
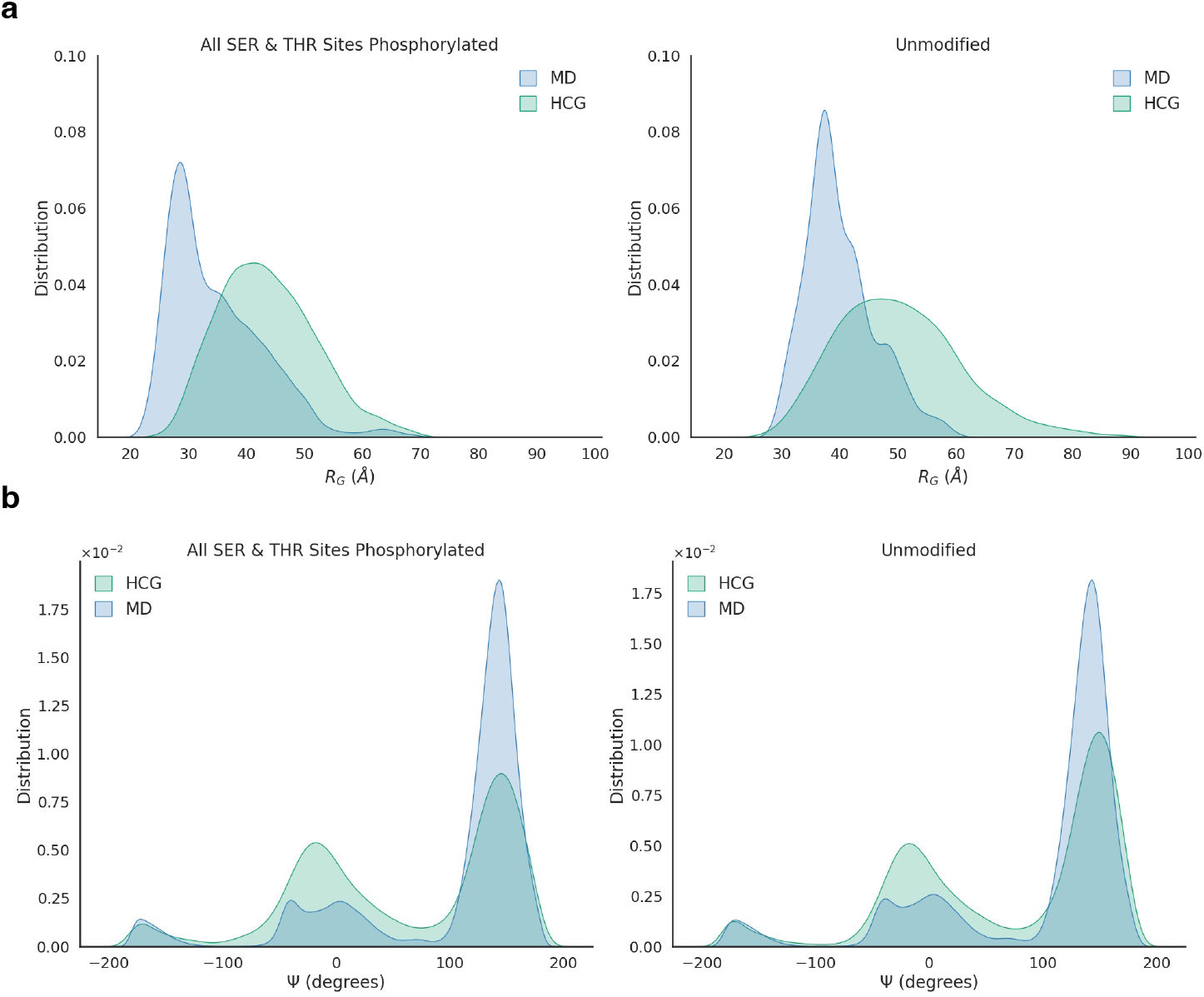
Evaluation of HCG for initial structural models. **(a)** Comparison of the radius of gyration (Rg) distributions from 1 µs MD simulations (blue) of the unmodified tau protein with CHARMM36m forcefield (right) and the modified peptide (left) against those generated by Hierarchical Chain Growth (HCG, green). **(b)** Similar comparison for the distribution of the ψ dihedral angle across residues in both the unmodified tau (right) and modified tau (left).

### Force Field Benchmarking and Validation

To determine the most suitable force field for the tau protein simulations, we compared AMBER03WS (with TIP4P-2005s water model)^18^, AMBER99SB*-ILDN-Q (with TIP4P-D)^19^, and CHARMM36m^20^ (with TIP3P-epsilon) using the experimentally well-characterized tau-K18 fragment. All force fields captured extended conformational distributions consistent with SAXS data reported by Mylonas et al.^21^ Among these, CHARMM36m exhibited superior sampling of extended conformations and better agreement with experimental residual dipolar coupling (RDC) data, making it the best choice for all subsequent simulations. Predicted RDCs were computed using PALES ^22^, following a previously established protocol^17^, and compared against experimental datasets ^22^**(Figure 3a-c**).

**Figure 3:**
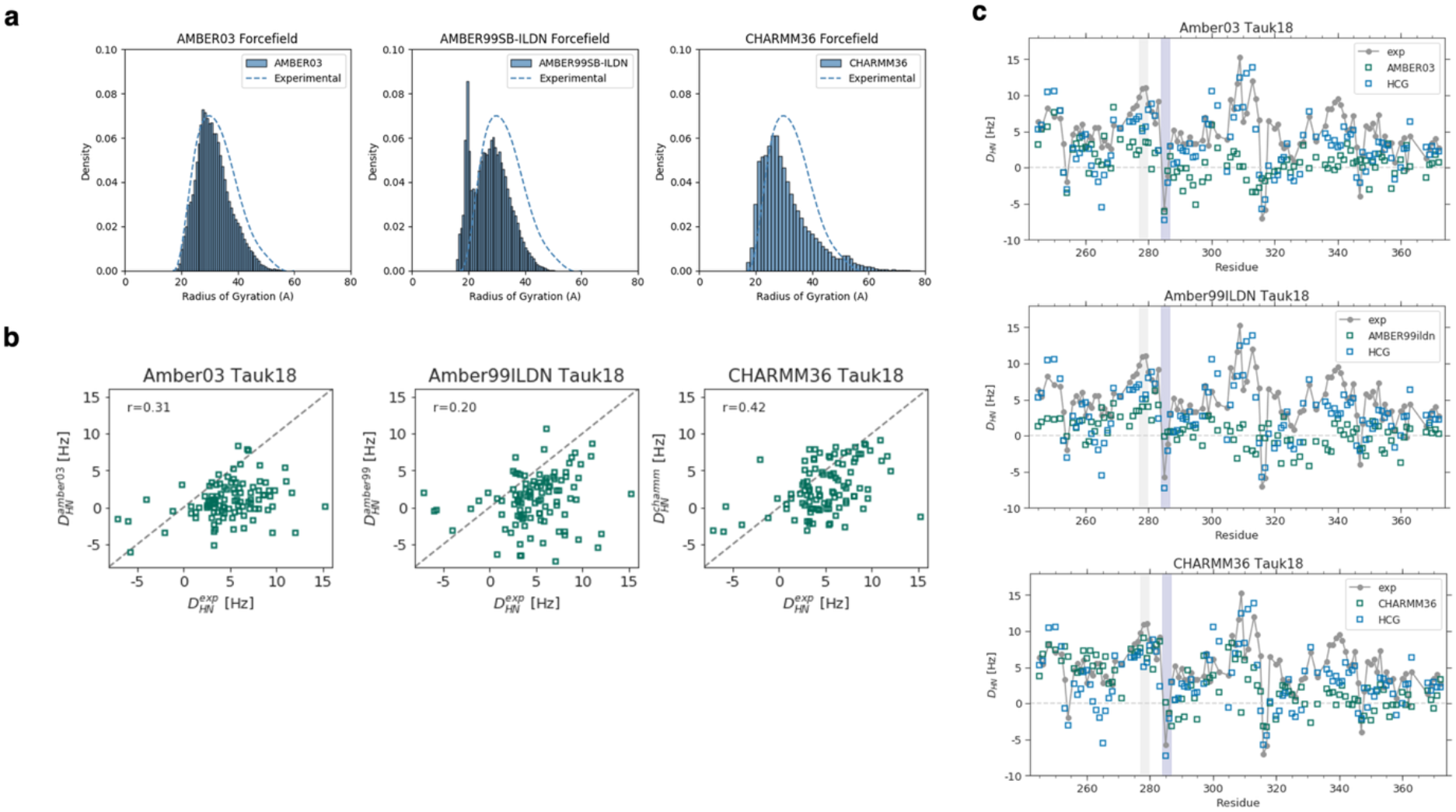
Validation of force fields. **(a)** Radius of gyration distributions for tau-K18 from 1µs (5 replicas) simulations using three different force fields (left: AMBER03, middle: Amber99SB*-ILDN-Q, right: CHARMM36m), compared to SAXS experimental data ^21^(dashed lines). **(b)** Scatter plot of calculated from MD output against measured experimental RDCs ^40^ for each forcefield. **(c)** Comparison of experimental and calculated ^1^H–^15^N RDCs from MD simulation output (1µs). Also shown are the RDCs from the HCG starting structures built from fragment MD simulations with the Amber99SB*-ILDN-Q force field.

### PTM Effects on Tau Condensates

To investigate the effects of PTMs in higher-order assemblies, we simulated condensates comprising 60 chains of the respective tau variant using the CHARMM36m force field for 1–2 µs per system (**Figure 4**). Fully modified chains yielded significantly more compact condensates, whereas single- and double-site modifications had minimal impact on compaction **(Figure 5a)**. Interestingly, hyper-modified chains formed fewer contacts with other chains, showing more compact, partially globular conformations **(Figure 5b)**. In parallel, self-interactions increased within a chain for modified residues. Tau remained largely disordered across condensate conditions, but specific PTMs modestly shifted its structure: phosphorylation at S202/T205 and acetylation at K280 increased the β-sheet content by ∼6–8%, full acetylation promoted helical structure (+3.5%), and full phosphorylation produced balanced increases in helices (2.8%) and sheets (4.7%) **(Figure 6 and Table S2)**. Notably, both fully acetylated and fully phosphorylated tau undergo near-immediate collapse within the first 0–200 ns of condensate simulations **(Figure S1)**, highlighting the strong impact of cumulative modifications on global chain compaction.

**Figure 4.**
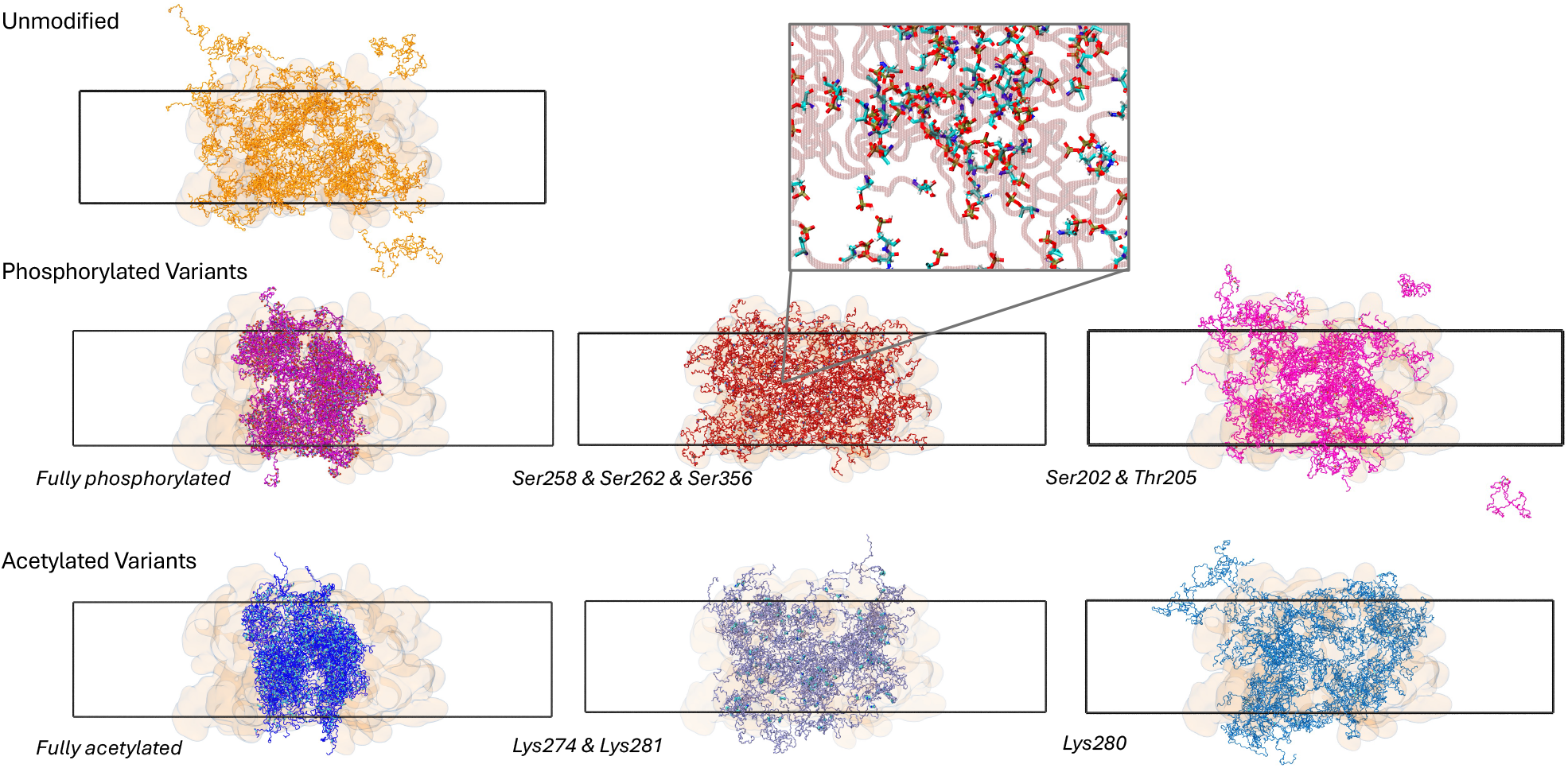
Effect of PTMs on condensates. **(Upper left)** Snapshots of the unmodified tau-K32 condensate in its initial conformation (orange transparent surface representation) and after 1.6µs of MD simulation. The MD simulations all started from the same equilibrated unmodified starting structure), and were left to run for 1.6 µs each in 3 replicas. The middle row shows the phosphorylated systems: Fully phosphorated, Ser258/Ser262/Ser356, Ser202/Thr205 (left to right). The bottom row shows the acetylated systems: Fully acetylated, Lys274/Lys281, and Lys280 (left to right). The dark lines indicate the boundaries of the simulation box.

**Figure 5:**
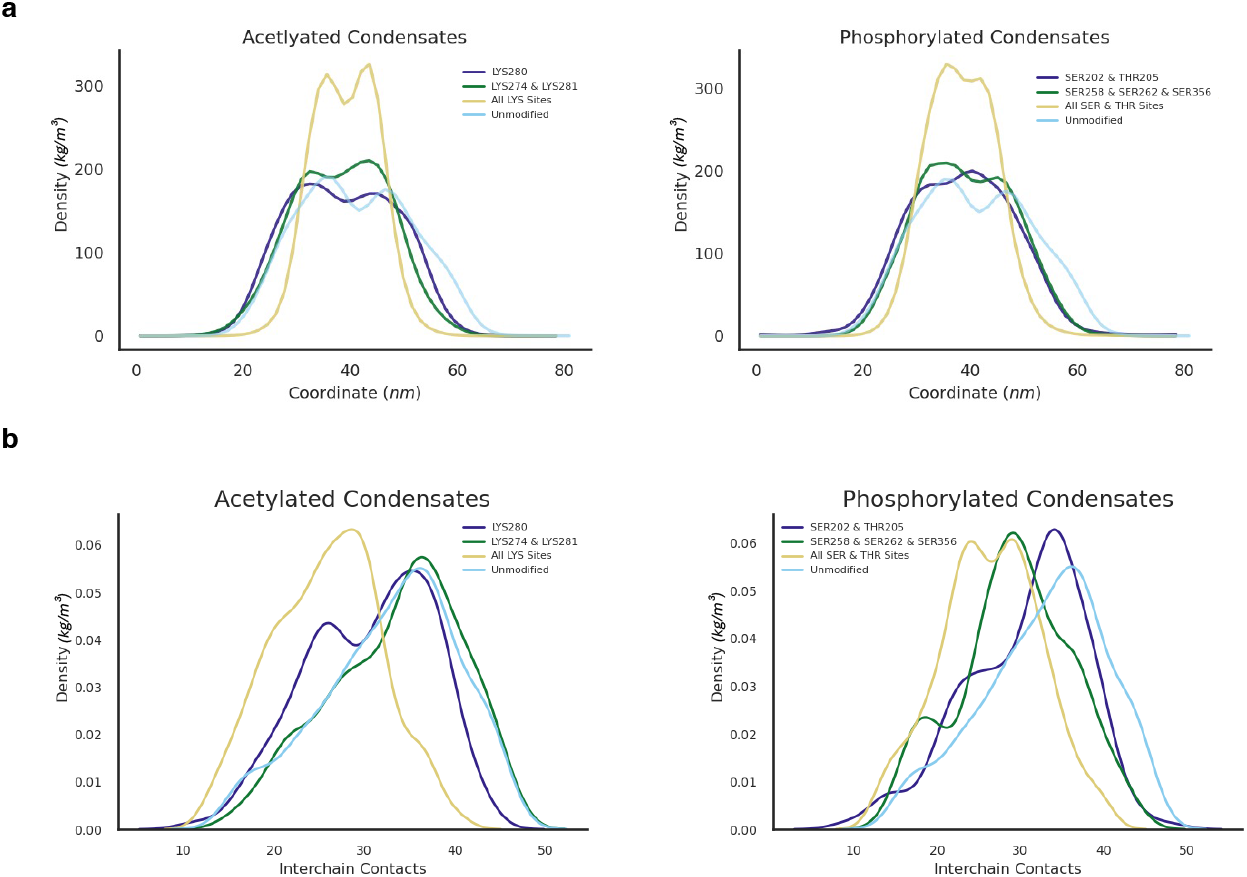
Effect of PTMs on condensate density profiles. **(a)** Protein density profiles for the condensate systems. Results are averaged over 3 replicas and shown for acetylated (left) and phosphorylated (right) chains of tau-K32. **(b)** Normalized probability density for the number of interchain amino-acid contacts in the condensate MD simulations (with a C_α_–C_α_ distance cutoff of 0.8 nm).

**Figure 6:**
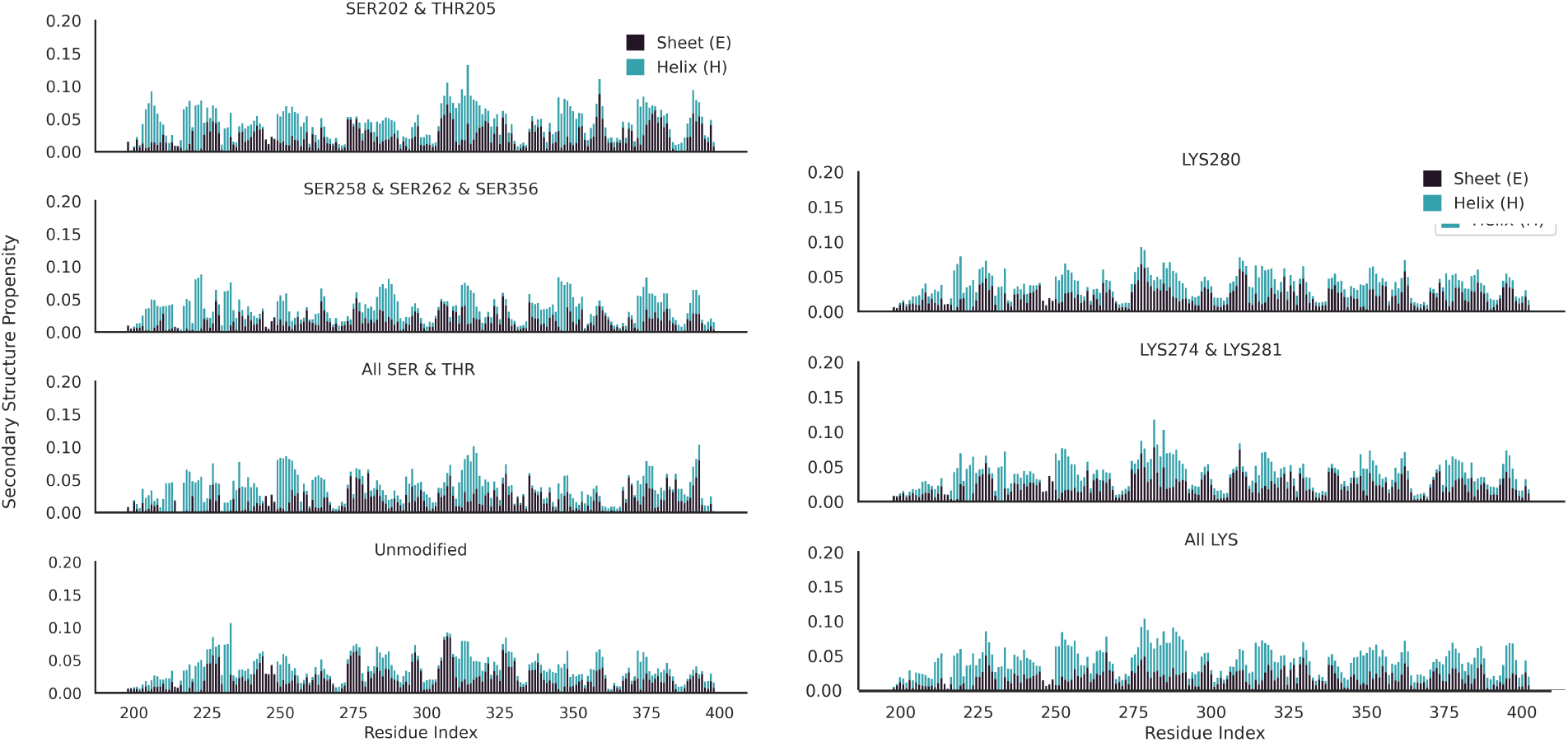
Time-averaged secondary structure distribution across all chains for each modified and unmodified tau in the condensate systems. This compares the distribution of helices (‘H’), strands (‘E’), and coils (‘C’) per residue in the chains. This is further described in **SI Table 2**.

Single-, double, and triple-site modifications showed more nuanced effects compared to the dramatic collapse of fully modified chains. Acetylation at Lys280 alone led to slightly more compact condensates compared to unmodified tau, while the Lys274/Lys281 pair produced a more noticeable though still modest reduction in compaction, consistent with these residues’ positioning in aggregation-prone motifs where acetylation can alter intermolecular contacts. Similarly, phosphorylation at S202/T205 and S256/S262/S365 led to modestly more compact condensates. These findings suggest that while isolated modifications can tune condensate properties in residue-specific ways, broad hyper-modification is required to trigger the pronounced collapse and self-compaction seen in fully modified tau. Interestingly, and in contrast to all other modifications we investigated, acetylation at Lys274/Lys281 did not reduce the number of interacting chains, but retained the distribution also observed for the unmodified state (**Figure 5**). This unexpected condensate behavior motivated a deeper investigation into the intrinsic effects of PTMs on isolated tau chains.

### Single-Chain PTM Effects on Tau Conformation

To disentangle intrinsic conformational effects from condensate crowding, we conducted 1–2 µs simulations for each of the seven systems using GROMACS and CHARMM36m, each with five replicas. Fully modified tau-K32 chains—either phosphorylated or acetylated—adopted more compact conformations. This effect was expected and arises from charge modulation: unmodified tau-K32 is highly positively charged, whereas acetylation neutralizes lysine side chains by removing positive charges, and phosphorylation introduces additional negative charges (see **Figure 1**). In both cases, the modifications shift tau toward a more balanced overall charge distribution, driving compaction (**Figure 7a & 7b**). More intriguingly, already only a few modifications had substantial effects in changing global conformations, especially acetylations at i) Lys280 and ii) Lys274 and Lys281 (**Figure 8a & 8b**).

**Figure 7:**
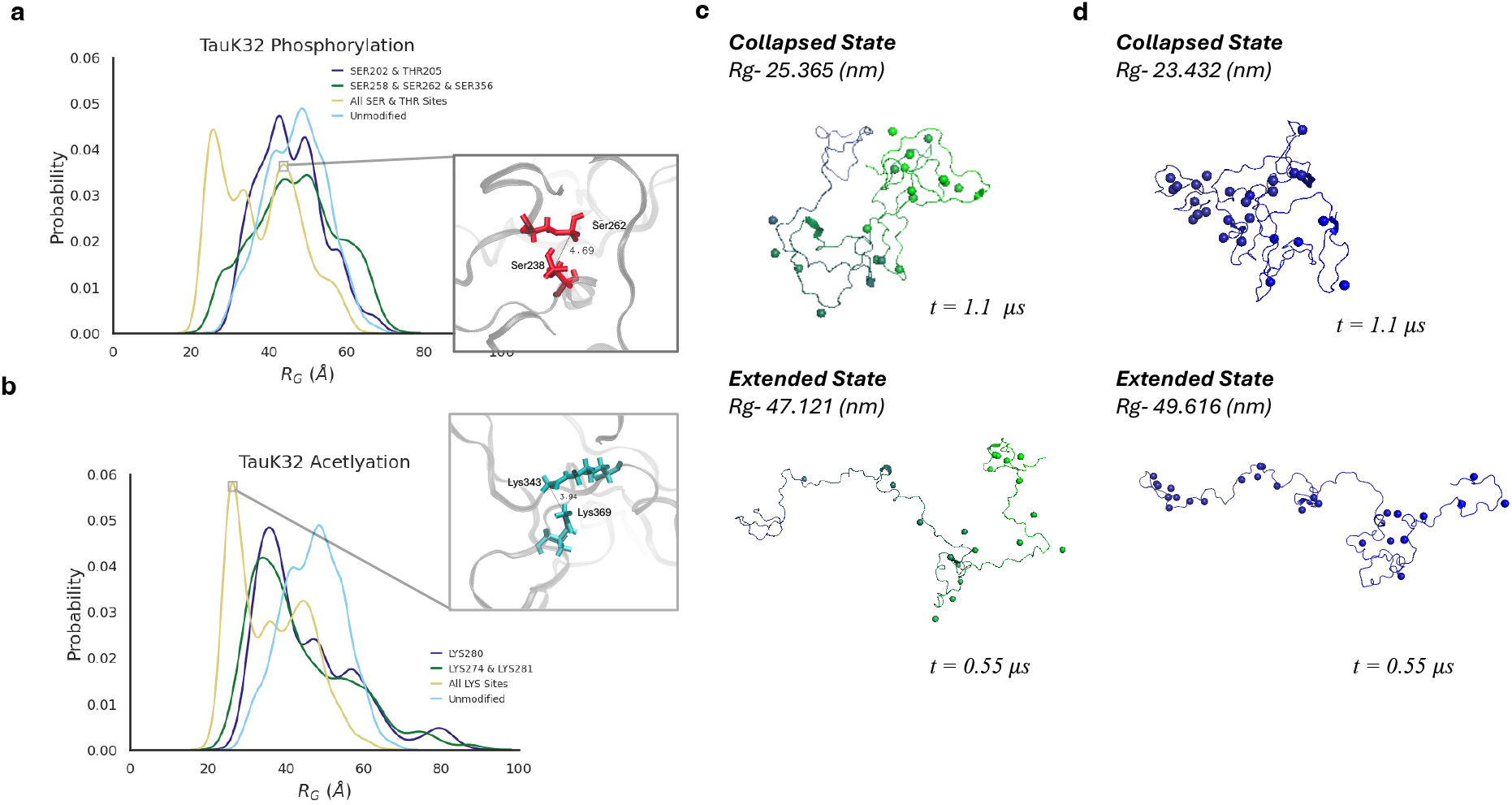
Single-chain tau-K32 protein conformational states – extended vs. collapsed. **(a)** Radius of gyration distributions for phosphorylated tau in three conditions: i) Ser202 & Thr205; ii) Ser258, Ser262 & Ser356; iii) and fully phosphorylated tau, overlaid with the unmodified tau chain. Representative simulation snapshot showing phosphoserines Ser238 and Ser262 in close proximity (P–P distance 4.69 Å), illustrating a hydrogen-bond interaction between phosphate groups that promote local chain collapse.**(b)** Average radius of gyration distributions for acetylated tau in three conditions, acetylation at: i) Lys280, ii) Lys274/Lys281, and iii) fully acetylated, also overlaid with unmodified tau. Representative snapshot of acetyl-lysines Lys343 and Lys369 (3.94 Å apart), showing the stacking of the carbonyl groups that stabilizes compacted conformations. **(c, d)** Example snapshots showing extended and collapsed conformations extracted at two different time points from the ∼1 µs simulations of fully phosphorylated **(c)** and fully acetylated **(d)** tau-K32.

**Figure 8:**
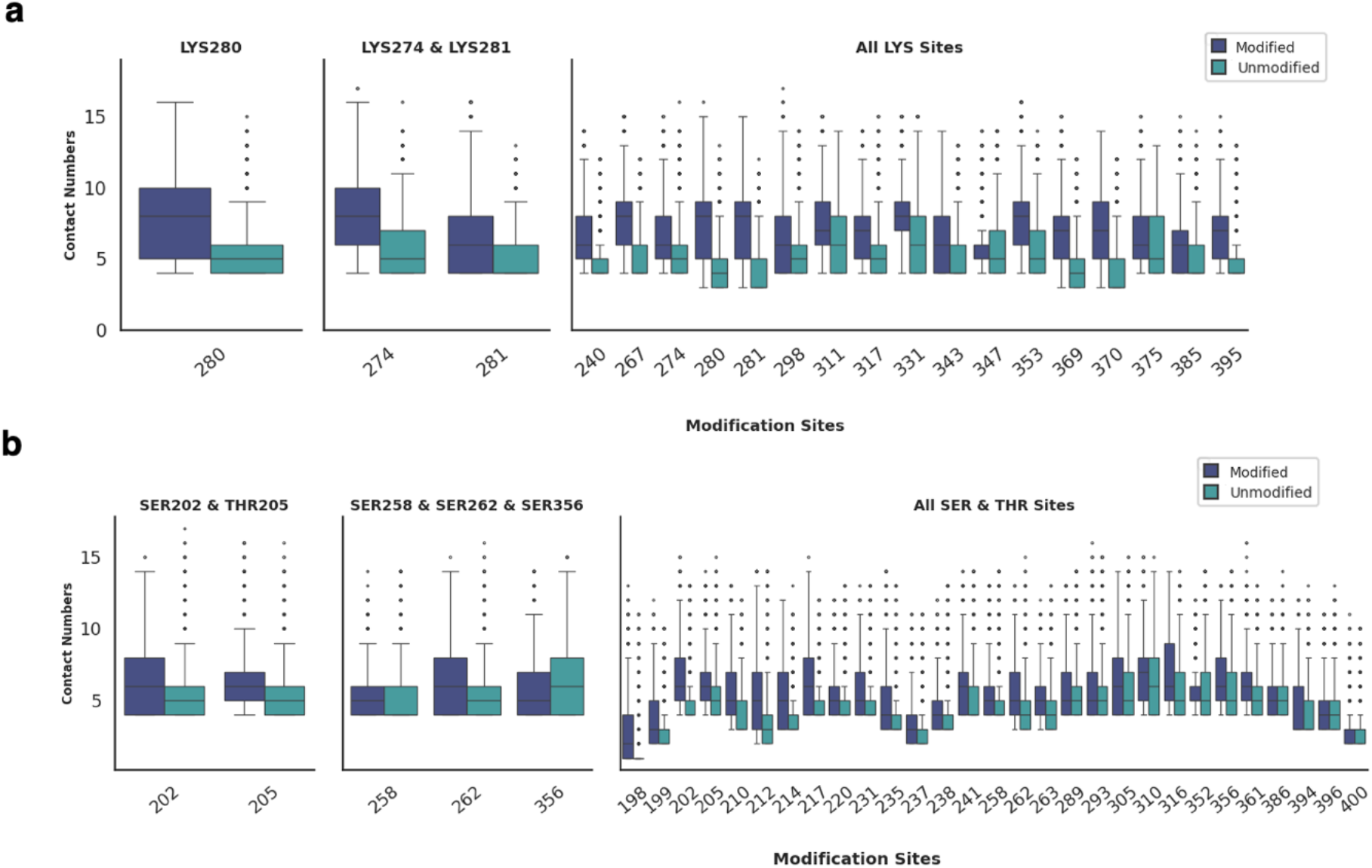
Contact frequency at modification sites in single-chain tau. Boxplots show the number of Cα–Cα contacts (<0.8 nm distance) between modified residues and the rest of the protein across MD simulations, comparing acetylated and unmodified systems (a), and phosphorylated and unmodified systems (b). On average, acetylated lysine sites show slightly higher contact numbers than unmodified (e.g., Lys280: 8.1 ± 3.0 vs. 5.3 ± 1.5; Lys274/Lys281: 7.4 ± 2.8 vs. 5.4 ± 1.7; all LYS sites combined: 6.9 ± 2.3 vs. 5.1 ± 1.8). Similarly, phosphorylated serine/threonine sites display modest increases relative to unmodified (e.g., Ser202/Thr205: 6.3 ± 2.1 vs. 5.2 ± 1.6; Ser258/Ser262/Ser356: 5.9 ± 1.9 vs. 5.5 ± 1.8; all SER/THR sites combined: 5.3 ± 2.2 vs. 4.5 ± 1.9). The minimum contact number of 4 (lower for terminal residues) is caused by the two residues before and after each residue always being in range of the cutoff.

Fully modified systems exhibited bimodal distributions of the radius of gyration (*Rg*), with two local minima near 26 Å and 43 Å (**Figure 7c &7d**,) indicating coexistence of compact and extended conformational states. We defined conformations with Rg below 30 Å as collapsed, and those above as extended. To identify structural features stabilizing each state, we compared residue-residue contacts between the collapsed and extended ensembles **(Figure S3a-d)**. In the fully phosphorylated system, key residues - Thr231, Arg230, Val229, Leu376, Lys375, and His374 - were associated with compaction. Similarly, in the fully acetylated system, residues Gly204, Pro203, Aly353, Ser352, and Gln351 played a stabilizing role **(Figure S3)**. Importantly, these changes extended beyond the immediate PTM sites, suggesting that modifications triggered broader conformational rearrangements across the tau chain. Residue-residue contact analysis revealed that PTMs enhanced interactions of modified residues **(Figure 7)**, while the overall protein remained globally disordered. Again, for the lightly modified systems, stronger effects could be observed for acetylation as compared to phosphorylation. The impact of individual modifications varied substantially across our simulations. Some sites showed little to no effect on structural metrics, while others produced clear shifts in compaction and contact patterns. In several cases, the apparent magnitude of the effect depended on the particular replica or sampling window, raising the question of whether these differences arise from limited sampling or reflect genuine underlying heterogeneity.

### Secondary Structure Propensity

Analysis of secondary structure content confirmed that tau remains predominantly disordered under all conditions. Coils dominated (>0.9 across systems), with only minor contributions from helices (0.02 ± 0.03) and β-sheets (0.04) in the unmodified state **(Figure 9)**. PTMs induced modest but distinct shifts: acetylation at Lys280 increased β-sheets by ∼7.6% percentage points, while full acetylation raised helical content by ∼3.5%. Phosphorylation at Ser202/Thr205 enhanced β-structure (∼5.9%), and full phosphorylation produced balanced gains in helices (∼2.8%) and β-sheets (∼4.7%) **(Table S3)**. These subtle, context-dependent changes may bias tau toward aggregation-prone or transient intermediates. Residue-residue interaction heatmaps revealed that modifications enhance local contacts at the modified sites while also driving broader rearrangements across the sequence (**Figure 10a**).

**Figure 9:**
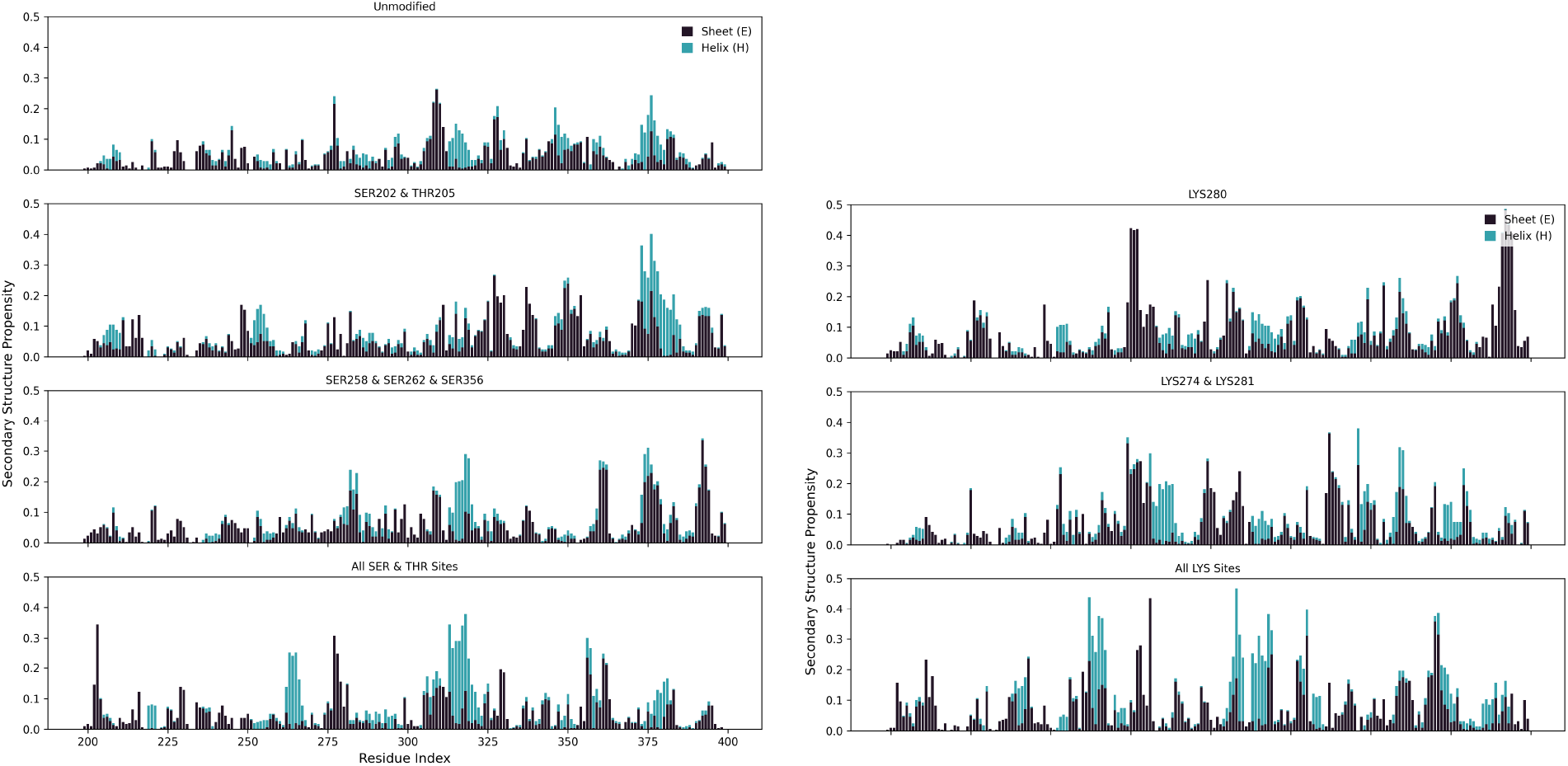
Time-averaged secondary structure distribution across the sequence for each modified and unmodified tau-K32 system. Stacked bar plots show per-residue propensities for coil, helix, and sheet structures, averaged over MD simulations. Each subplot represents a different phosphorylation, acetylation system, or the unmodified tau system.

**Figure 10:**
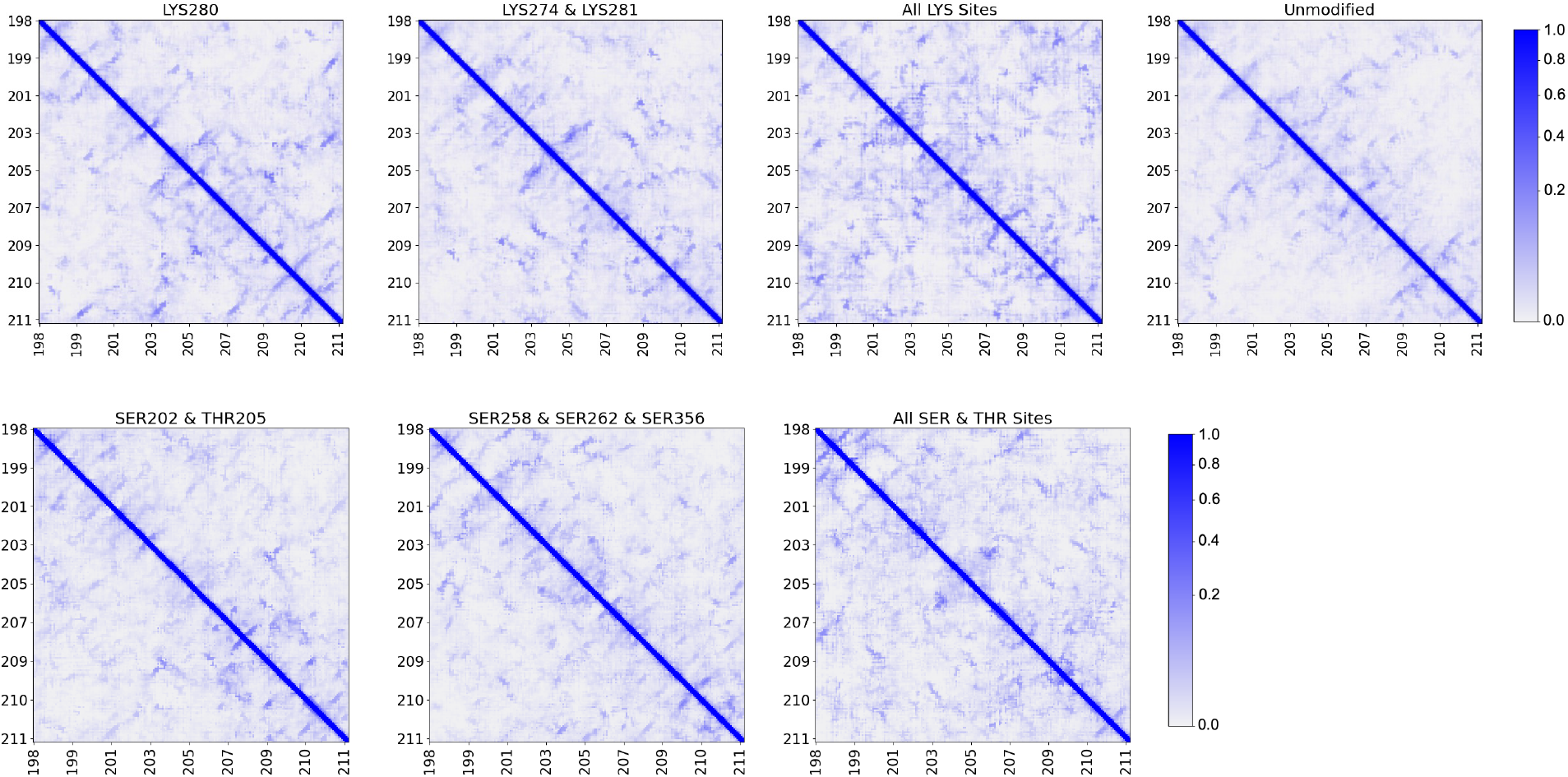
Protein–protein interaction maps for modified and unmodified tau. Normalized contact frequency heatmaps computed using a 0.5 nm cutoff between all closest-heavy atoms across simulation trajectories. Each panel corresponds to a different modification state. Higher intensity indicates more frequent residue–residue contacts over time.

### Limitations

All-atom MD simulations of condensates are challenging for multiple reasons, in particular the quality of force fields, the slow relaxation times of conformational states in dense protein solutions, and the even slower relaxation of the equilibrium between dilute and dense phases. Here, these challenges are compounded by the presence of PTMs, in particular phosphorylation, which can further enhance interactions and thus slow down dynamics^23,24^. As a result, our condensate simulations—and to a lesser degree also the single tau-K32 simulations—are likely to be representative but not fully converged.

To address the known problems of developing balanced force fields for disordered proteins, we screened different candidates against experimental data for a shorter tau fragment. While this should address the major issues, the problem of accurate PTM modelling remains. For the phosphorylation, doubly negative charge states tend to lead to unphysiological aggregation^25^. To avoid these problems, we settled on the singly negative charge state of the phosphate group, even though at physiological conditions we would expect to have substantial population of both states.

Finally, we noticed in the post-analysis that the soft-core minimization used for the condensate systems had introduced some chirality errors into the peptide sequences in the condensate systems. Specifically, the stereochemistry of a small number of residues was altered due to inversion of the chiral center to the incorrect configuration^26,27^. We do not expect these errors influenced the outcomes of our simulations, and the affected residues are reported in Supporting Information **Table 6**.

### Conclusions

This study systematically examined the structural and aggregation-related consequences of site-specific and cumulative PTMs on the tau-K32 fragment using atomistic MD simulations. Employing HCG-derived starting structures and the CHARMM36m force field, we evaluated seven tau-K32 variants with experimentally validated PTM patterns.

Our results show that tau-K32 remains largely disordered across all conditions, yet single-point modifications can exert surprisingly strong effects on global chain properties. For instance, acetylation at Lys280 or phosphorylation at Ser202/Thr205 introduced localized biases toward β-sheet content that propagate to influence overall conformation. Hyper-modification drives near-immediate chain collapse, particularly notable for acetylation: while not introducing new sites for salt bridge interactions – as phosphorylation does – acetylation also reduces net positive charge, yielding a more neutral condensate, analogous to phosphorylation, which adds negative charge to achieve charge balance.

Full phosphorylation and acetylation increased conformational heterogeneity, with sampling of both compact and extended states. In condensate simulations, fully modified systems form more compact assemblies, consistent with single-chain behaviour. These findings highlight how PTMs modulate both local residue interactions and global chain architecture, providing mechanistic insight into tau’s aggregation propensity and offering a framework to guide therapeutic strategies for tauopathies.

## Materials and Methods

### Generation of HCG-Derived Tau Structures

To construct structural ensembles for the tau-K32 fragment (UniProt ID: P10636-8), we used the HCG algorithm developed by Pietrek et al.^16^. The protein sequence was divided into six overlapping fragments spanning the proline-rich and C-terminal domains, excluding the N1 and N2 inserts. Overlapping residues were used to enable proper stitching of neighbouring fragments. Each fragment was subjected to 20 μs of replica-exchange molecular dynamics (REMD) simulations using the Amber99SB*-ILDN-Q force field with the TIP4P-D water model. Simulations were conducted with 24 replicas across a temperature range of 288–431 K. The resulting fragments were assembled into 1,000 full-length structures using the HCG algorithm, which were then used as initial configurations for MD simulations. The fragments were defined as follows:

**P2** - SSPGSPGTPGSRSRTPSLPTPPTREPKKVAVVRTPPKSPSSAKSRL**QT**

**R1**-**QT**APVPMPDLKNVKSKIGSTENLKHQPGGGK**VQ**

**R2**-**VQ**IINKKLDLSNVQSKCGSKDNIKHVPGGGS**VQ**

**R3**-**VQ**IVYKPVDLSKVTSKCGSLGNIHHKPGGGQ**VE**

**R4**-**VE**VKSEKLDFKDRVQSKIGSLDNITHVPGGGN**KK**

**R**-**KK**IETHKLTFRENAKAKTDHGAEIVYKSPVVS

We chose longer fragments than normally used in HCG (dimers or pentamers) to capture at least some of the more long-range electrostatic interactions.

### Force Field Benchmarking

Force field validation was performed on the tau-K18 fragment, for which experimental SAXS and RDC data are available. We evaluated AMBER03WS, AMBER99SB*-ILDN-Q ^19^, and CHARMM36-July2021^28,29^ (in the text noted as CHARMM36m in short), each with corresponding water models (tip4p2005s, tip4pd, tip3p-epsilon; respectively). Simulations were conducted using GROMACS 2021.5^1^ and solvated in a dodecahedral box with a buffer of 2.5 nm on either side at a 150 mM NaCl concentration. The solvated system was then energy minimized with the steepest descent algorithm (target maximum force of 500 kJ/mol/nm) with a step size of 0.01 nm, and canonical ensemble equilibration (NVT) was carried out for 50000 steps at a 2 fs timestep at 300 K with a characteristic time of 0.1 ps velocity rescaling^32^, and isothermal-isobaric equilibration (NPT) was done for 125000 steps (2 fs timestep) at 300 K using the Berendsen barostat^33^ (1.0 ps time constant and reference pressure of 1.0 bar). Production runs for all 10 replicas were carried out for 1.1–1.6 µs simulations at 300 K (2fst timestep and velocity rescaling temperature coupling, 0.1ps time constant; and Panrinello-Rahman pressure coupling with 1.0ps time constant and reference pressure of 1.0 bar). All equilibration and production steps were done with LINCS constraints on all bonds with hydrogens, VDW interactions cutoff at 1.2 nm, full periodic boundary conditions, and a particle-mesh Ewald for long range electrostatic interactions. CHARMM36-July2021 was selected based on its superior performance in capturing extended conformations, matching RDC values, and supporting PTM parameters.

### Simulation Protocols and Post-Processing

PTMs were introduced using CHARMM-GUI^34^, and simulations were conducted using GROMACS 2021.5 with the CHARMM36-July2021 force field and TIP3P-ε water. The systems were all built the same as above, and each of the seven tau systems was simulated for 1–2 µs with five independent replicas.

Structural and conformational analyses were performed using GROMACS built-in tools and the MDAnalysis tools tools^35,36^. Residue-residue contact maps were computed as the minimum heavy-atom distance between each pair of residues across all frames, with a contact defined by a cutoff of 1.2 nm. Mass density profiles along the simulation box axis were computed using the gmx density command, which outputs density in kg/m^3^, and were used to assess the spatial distribution of the chain and solvent. Secondary structure content was assigned for every frame using the DSSP algorithm as implemented in MDAnalysis, which classifies each residue into helix, beta-strand, turn, or coil states based on backbone hydrogen bond geometry. Fractional secondary structure populations were computed by averaging assignments over all frames and replicas. Back-calculated ^1^H-^1^5N residual dipolar couplings (RDCs) were computed using the PALES software^22^. PALES predicts the molecular alignment tensor from the three-dimensional structure of each snapshot using steric obstruction modeling, then uses this tensor to compute the expected RDC for each amide bond vector. Back-calculated RDCs were averaged over the simulation ensemble and compared to experimental values to assess force field performance. Molecular visualizations and trajectory snapshots were generated using VMD and PyMOL^37,38^.

### Building and Running Condensate CG Systems

An atomistic starting structure was coarse-grained using martinize to produce a Martini 2.2 coarse-grained (CG) representation of a single tau-K32 chain. The single CG chain was first equilibrated in a cubic box to obtain a collapsed conformation representative of the disordered chain ensemble.

The collapsed CG chain was then assembled into a 60-chain condensate configuration using PyMOL, placed in a cubic box (150 × 150 × 150 Å initial dimensions prior to solvation). The condensate was built using the Martini 2.2 force field with scaled protein-protein interaction parameters and GROMACS 2020.6. The system was solvated with Martini water beads (W) and antifreeze water beads (WF, ∼10% fraction) using a CG bead radius of 0.21 nm, neutralised, and brought to a physiological salt concentration of 150 mM NaCl, giving a final system of 60 tau-K32 chains and 267,257 CG particles total. Energy minimisation was performed twice using the steepest-descent algorithm, first in vacuum and then after solvation. NVT equilibration was performed using the velocity-rescaling thermostat at 300 K (100,000 steps, dt = 0.015 ps, 1.5 ns total, τT = 1.0 ps), followed by NPT equilibration using the Berendsen barostat at 1.0 bar with isotropic pressure coupling (500,000 steps, dt = 0.015, 7.5 ns total). This yielded an equilibrated cubic condensate box of approximately 203 × 203 × 203 Å.

The cubic condensate box was then extended along the X-axis to produce a slab geometry (approximately 827 × 200 × 200 Å), placing the condensate in the centre of an elongated simulation cell with dilute-phase solvent on either side. This slab geometry allows the condensate– dilute phase interface to form and equilibrate naturally under periodic boundary conditions.

Production simulations were run at an intermediate-to-high protein–protein interaction scaling factor of α = 0.75, using the Parrinello-Rahman barostat with anisotropic pressure coupling applied exclusively along the X-axis (compressibility = 3 × 10^−4^ bar^−1^ in X, zero in Y and Z), maintaining the slab geometry throughout. All simulations used the Verlet neighbour-list scheme, a cutoff of 1.1 nm for both Coulomb and van der Waals interactions with potential-shift modifiers, and a dielectric constant of εr = 15. Production runs were performed for 30 µs at a timestep of 30 fs.

For downstream atomistic analysis, CG condensate snapshots were back-mapped to full atomistic resolution using the CG2AT software^39^, selecting the de novo backmapping protocol. Three independent atomistic replicas were generated from the last three trajectory frames of the converged Martini simulation, which were then used as starting configurations for atomistic MD simulations. Due to the size of the condensates, PTMs could not be introduced manually via CHARMM-GUI. Instead, a custom Python script was employed to determine the position of a target atom (D) using three reference atoms (A, B, and C), along with bond length, bond angle, and dihedral angle values obtained from the force field. The script projected atom A onto the B–C plane and solved geometric constraints to position atom D, allowing for the insertion of acetyl or phosphate groups.

### Atomistic Condensate Simulations

Atomistic condensate simulations were performed using GROMACS 2022.5 with the CHARMM36-July2021 force field and TIP3P-ε water, following a similar protocol as the single-chain simulations except where noted below. The three backmapped condensate replicas (∼189,780 atoms, rectangular box ∼83×20×20 nm) served as starting structures. Prior to equilibration, each system was solvated in TIP3P-ε water and 150 mM NaCl was added (2770 Na^+^ and 3910 Cl^−^ ions, including counterions to neutralize the net charge). Due to the structural complexity of the backmapped condensate, a soft-core pre-equilibration protocol was required to relieve steric clashes prior to standard equilibration. This consisted of three sequential steepest-descent energy minimizations (up to 5,000 steps each, emstep=0.01) using progressively harder soft-core potentials within the GROMACS free-energy framework (λ=0.01, sc-power=2, couple-intramol=yes; sc-α=4, then 1,then 0.1), followed by a standard steepest-descent energy minimization (up to 50,000 steps, emtol=1000 kJ/mol/nm, PME electrostatics, rcoulomb=rvdw=1.2 nm). NVT equilibration was carried out for 50,000 steps at dt=2 fs (100 ps) at 300 K using velocity rescaling (τT=0.1 ps) with position restraints. NPT equilibration was performed in two stages: first with position restraints (125,000 steps, dt=2, fs = 250 ps, Berendsen barostat, anisotropic, τP=2.0 ps, 1.0 bar), then without position restraints using the same parameters to allow full system relaxation. Production runs were carried out for 1.1–1.5 μs at dt=2 fs, across 3 independent replicas, using the Parrinello-Rahman barostat (anisotropic, τP=2.0 ps, 1.0 bar) and V-rescale thermostat (τT=0.1 ps, 300 K).

## Acknowledgements

We thank Maziar Heidari for help with the setup of the condensate systems. We thank the Max Planck Computing and Data Facility for computational resources and the Max Planck Society for financial support.

## Notes

### Competing Interest Statement

The authors have declared no competing interest.

