## Supplemental Information for "Effects of PTMs on Tau Protein Aggregation: Insights from HCG and Atomistic MD Simulations"

### Supplementary Information

| System | Modification Sites |
|---|---|
| Fully Phosphorylated | S198,S199,S202,T205,S210,T212,S214,T217,T220,T231,S235,S237,S238,S241,S258,S262,T263,S289,S293,S305,Y310,S316,S352,S356,T361,T386,Y394,S396,S400 |
| Ser 202 & Thr 205 | S202/T205 |
| Ser 258 & Ser 262 & Ser 356 | S258/262/356 |
| Fully Acetylated | K240,K267,K274,K280,K281,K298,K311,K317,K331,K343,K347,K353,K375,K370,K369,K385,K395 |
| Lys 280 | K280 |
| Lys 274 & Lys 281 | K274,281 |

**Table 1. Tau Modification sites.**

| System | Modification Site | Coil (%) | Helix (%) | Sheet (%) |
|---|---|---|---|---|
| K280 | 280 | 96.4 ± 2.1 | 1.4 ± 1.4 | 2.2 ± 1.5 |
| K274, K281 | 274, 281 | 96.2 ± 2.3 | 1.5 ± 1.6 | 2.3 ± 1.6 |
| All LYS Sites | multiple (full acetylation) | 95.9 ± 2.5 | 2.2 ± 1.8 | 1.9 ± 1.5 |
| Unmodified | none | | | |
| S202, T205 | 202, 205 | 95.9 ± 2.3 | 1.7 ± 1.7 | 2.4 ± 1.7 |
| S258, S262, S356 | 258, 262, 365 | 96.4 ± 2.2 | 1.7 ± 1.7 | 2.0 ± 1.4 |
| All SER & THR Sites | multiple (full phosphorylation) | 96.5 ± 2.3 | 1.8 ± 1.8 | 1.7 ± 1.5 |

**Table 2. Secondary Structure Statistics for Tau Condensate**

| System | Fragment | Coil (%) | Helix (%) | Sheet (%) |
|---|---|---|---|---|
| K280 | frag_1 | 93.10 ± 1.00 | 2.00 ± 1.10 | 4.90 ± 0.20 |
| | frag_2 | 94.40 ± 0.30 | 1.40 ± 0.10 | 4.20 ± 0.20 |
| K274, K281 | frag_1 | 93.50 ± 0.40 | 2.00 ± 0.40 | 4.50 ± 0.20 |

| | | | | |
|---|---|---|---|---|
| | frag_2 | 95.20 ± 0.60 | 0.40 ± 0.10 | 4.40 ± 0.50 |
| All LYS Sites | frag_1 | 92.40 ± 0.60 | 4.20 ± 0.80 | 3.40 ± 0.20 |
| | frag_2 | 95.80 ± 0.90 | 1.90 ± 0.70 | 2.30 ± 0.50 |
| Unmodified | frag_1 | 95.30 ± 0.50 | 0.80 ± 0.10 | 3.90 ± 0.50 |
| | frag_2 | 93.80 ± 0.80 | 0.90 ± 0.30 | 5.30 ± 0.70 |
| S202, T205 | frag_1 | 94.80 ± 0.70 | 1.10 ± 0.30 | 4.10 ± 0.70 |
| | frag_2 | 92.90 ± 0.60 | 1.70 ± 0.60 | 5.40 ± 0.20 |
| S258, S262, S356 | frag_1 | 95.70 ± 0.80 | 1.30 ± 0.20 | 3.00 ± 0.60 |
| | frag_2 | 95.50 ± 0.50 | 0.90 ± 0.30 | 3.60 ± 0.20 |
| All SER & THR Sites | frag_1 | 94.90 ± 1.10 | 1.90 ± 1.20 | 3.20 ± 0.50 |
| | frag_2 | 96.40 ± 0.50 | 1.00 ± 0.20 | 2.70 ± 0.30 |

**Table 3. Secondary Structure Statistics for condensate Tau looking only at 2 aggregation-prone regions. Shows mean and SEM.**

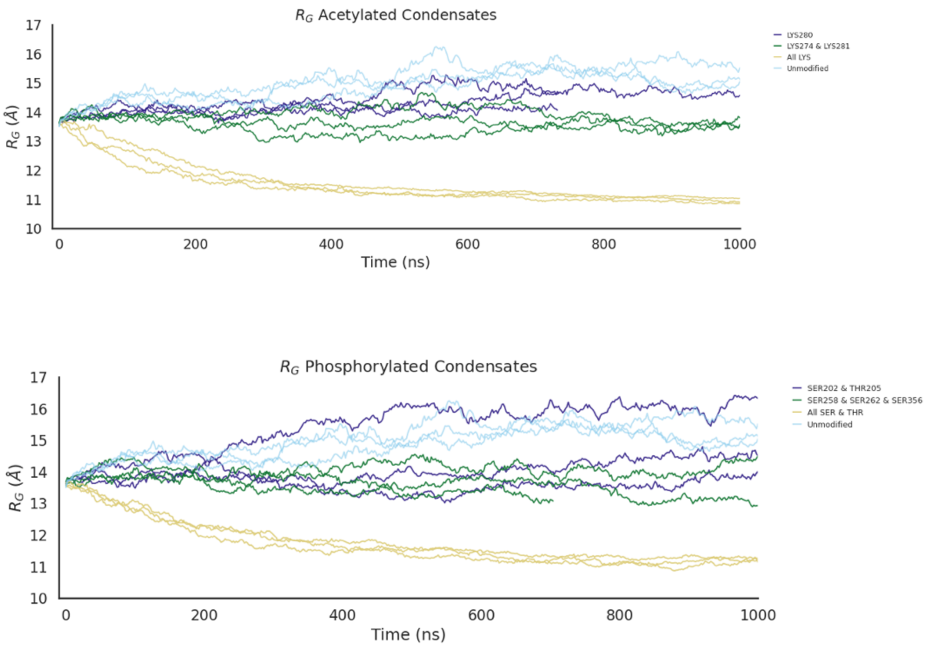


**Figure S1**: Time evolution of the radius of gyration for the overall condensate, shown for a single replica of acetylated and phosphorylated chains of Tau.

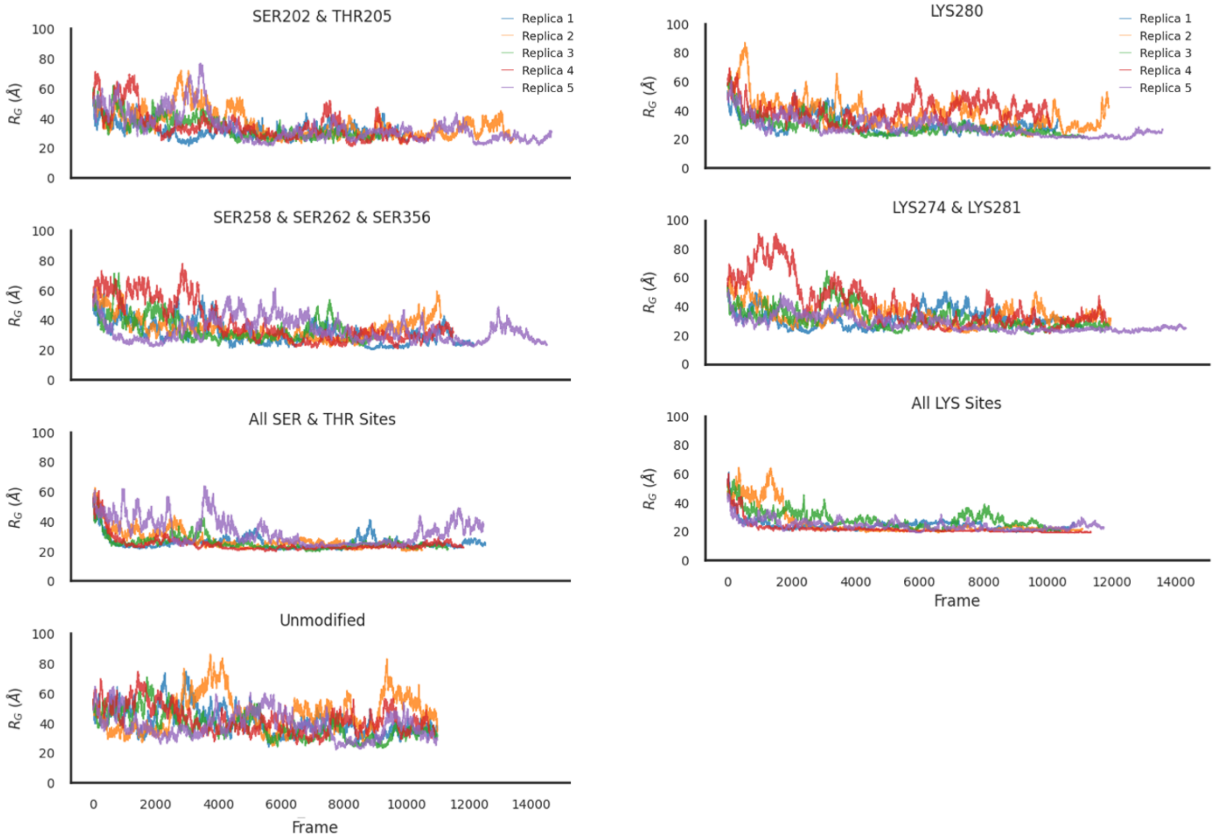


**Figure S2**: **Convergence analysis – single chain replicas**. Shows the radius of gyration over time for all 5 replicas in each system for single chain tau.

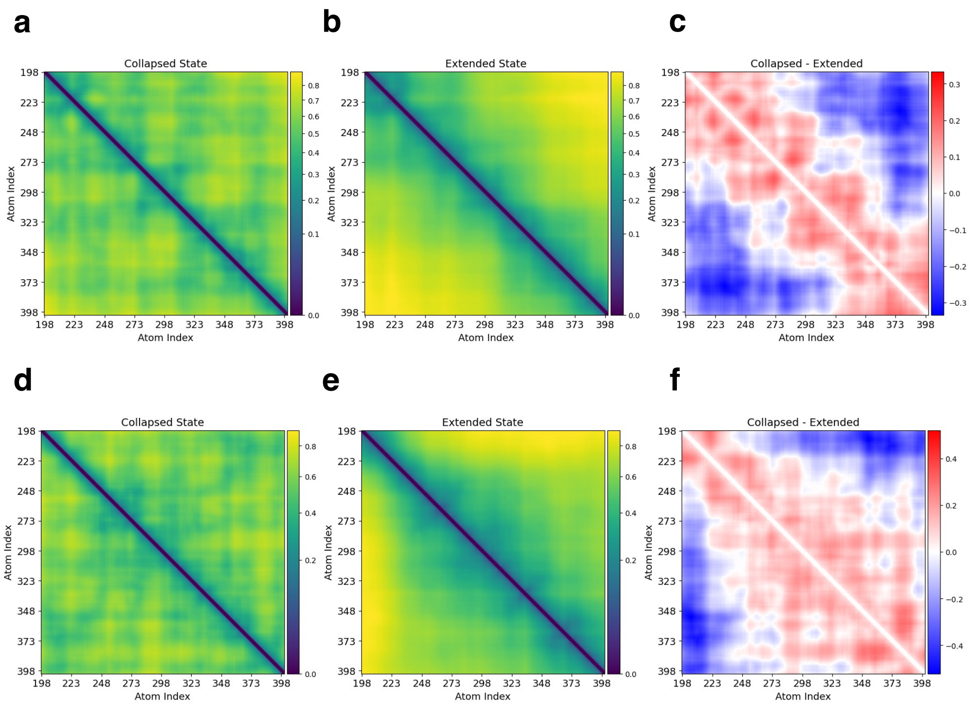


**Figure S3**. **Residue- residues Cα–Cα contacts (nm) for two states – collapsed and extended**. Heatmaps **(a)** and **(b)** represent the fully phosphorylated system in its two primary conformational states: Collapsed (lower minimum, $R_G$ <30 nm) and Extended (higher minimum, $R_G$ >30 nm), respectively. Panel **(c)** displays the difference in contact patterns between these two states. A similar analysis is shown for the fully acetylated system in panels **(d–f).**

| Modification Site | Coil (%) | Helix (%) | Sheet (%) |
|---|---|---|---|
| K280 | 90.96 ± 8.77 | 1.42 ± 1.84 | 7.63 ± 8.84 |
| K274, K281 | 91.66 ± 8.16 | 2.34 ± 3.77 | 5.99 ± 7.32 |
| All LYS Sites | 89.81 ± 10.1 | 3.47 ± 6.27 | 6.72 ± 7.45 |
| Unmodified | 94.27 ± 5.03 | 1.73 ± 2.75 | 4.00 ± 4.28 |
| S202, T205 | 91.96 ± 7.22 | 2.11 ± 4.08 | 5.93 ± 5.62 |
| S258, S262, S356 | 92.99 ± 7.85 | 1.64 ± 3.13 | 5.37 ± 5.56 |
| All SER & THR Sites | 92.53 ± 7.85 | 2.79 ± 5.73 | 4.68 ± 5.55 |

**Table 4. Secondary Structure Statistics for single chain Tau.** Shows mean and SEM.

| System | Fragment | Coil (%) | Helix (%) | Sheet (%) |
|---|---|---|---|---|
| K280 | frag_1 | 73.18 ± 6.48 | 0.06 ± 0.06 | 26.76 ± 6.51 |
| | frag_2 | 87.03 ± 3.10 | 0.57 ± 0.32 | 12.40 ± 3.33 |
| K274, K281 | frag_1 | 76.49 ± 13.45 | 0.99 ± 0.92 | 22.52 ± 13.82 |
| | frag_2 | 85.60 ± 4.83 | 0.15 ± 0.14 | 14.25 ± 4.90 |
| All LYS Sites | frag_1 | 85.21 ± 4.88 | 0.07 ± 0.04 | 14.72 ± 4.89 |
| | frag_2 | 75.93 ± 4.65 | 16.93 ± 7.28 | 7.14 ± 3.03 |

| | | | | |
|---|---|---|---|---|
| Unmodified | frag_1 | 91.13 ± 1.97 | 1.67 ± 1.31 | 7.20 ± 0.83 |
| | frag_2 | 82.30 ± 4.75 | 0.50 ± 0.41 | 17.21 ± 4.47 |
| S202, T205 | frag_1 | 94.20 ± 2.15 | 0.27 ± 0.14 | 5.53 ± 2.19 |
| | frag_2 | 90.39 ± 1.70 | 1.27 ± 0.82 | 8.34 ± 2.18 |
| S258, 262, 356 | frag_1 | 93.39 ± 2.73 | 1.50 ± 1.12 | 5.11 ± 2.55 |
| | frag_2 | 89.66 ± 2.51 | 0.81 ± 0.36 | 9.53 ± 2.26 |
| All SER & THR Sites | frag_1 | 84.46 ± 6.19 | 0.03 ± 0.02 | 15.51 ± 6.19 |
| | frag_2 | 0.8483 ± 0.0316 | 0.0278 ± 0.0273 | 0.1238 ± 0.0389 |

**Table 5. Secondary Structure Statistics for single chain Tau looking only at 2 aggregation-prone regions.** Shows mean and SEM.

| System | Residue | Position |
|---|---|---|
| S258, S262, S356 | SP1 | 60 |
| S258, S262, S356 | SP1 | 64 |
| S258, S262, S356 | PRO | 53 |
| S258, S262, S356 | LEU | 159 |
| S258, S262, S356 | ASN | 161 |
| S258, S262, S356 | ASN | 129 |
| S258, S262, S356 | VAL | 141 |
| S258, S262, S356 | LYS | 149 |
| S258, S262, S356 | LYS | 61 |
| S258, S262, S356 | THR | 65 |
| S258, S262, S356 | PRO | 34 |
| S258, S262, S356 | PRO | 35 |
| S258, S262, S356 | SP1 | 158 |
| S202, T205 | PRO | 53 |
| S202, T205 | ASN | 129 |
| S202, T205 | VAL | 141 |
| S202, T205 | LYS | 149 |
| S202, T205 | PRO | 34 |
| S202, T205 | PRO | 35 |
| K280 | PRO | 53 |
| K280 | ASN | 129 |
| K280 | LYS | 149 |
| K280 | PRO | 34 |
| K274, K281 | ALY | 83 |
| K274, K281 | PRO | 53 |
| K274, K281 | ASN | 129 |
| K274, K281 | LYS | 149 |
| K274, K281 | PRO | 34 |
| All SER & THR Sites | SP1 | 37 |
| All SER & THR Sites | SP1 | 40 |
| All SER & THR Sites | SP1 | 43 |
| All SER & THR Sites | SP1 | 60 |
| All SER & THR Sites | SP1 | 64 |

| | | |
|---|---|---|
| All SER & THR Sites | SP1 | 91 |
| All SER & THR Sites | SP1 | 107 |
| All SER & THR Sites | SP1 | 118 |
| All SER & THR Sites | SP1 | 154 |
| All SER & THR Sites | SP1 | 158 |
| All SER & THR Sites | SP1 | 198 |
| All SER & THR Sites | SP1 | 202 |
| All SER & THR Sites | LYS | 61 |
| All SER & THR Sites | LYS | 92 |
| All SER & THR Sites | LYS | 119 |
| All SER & THR Sites | LYS | 149 |
| All SER & THR Sites | LYS | 187 |
| All SER & THR Sites | LEU | 117 |
| All SER & THR Sites | LEU | 159 |
| All SER & THR Sites | VAL | 30 |
| All SER & THR Sites | VAL | 89 |
| All SER & THR Sites | VAL | 141 |
| All SER & THR Sites | ILE | 130 |
| All SER & THR Sites | THR | 65 |
| All SER & THR Sites | THR | 179 |
| All SER & THR Sites | GLN | 90 |
| All SER & THR Sites | ASN | 129 |
| All SER & THR Sites | CYS | 93 |
| All SER & THR Sites | THZ | 163 |
| All SER & THR Sites | THZ | 188 |
| All SER & THR Sites | PRO | 34 |
| All SER & THR Sites | PRO | 38 |
| All SER & THR Sites | PRO | 53 |
| All SER & THR Sites | PRO | 103 |
| All LYS Sites | ALY | 82 |
| All LYS Sites | ALY | 83 |
| All LYS Sites | ALY | 100 |
| All LYS Sites | ALY | 113 |
| All LYS Sites | ALY | 133 |
| All LYS Sites | ALY | 145 |
| All LYS Sites | ALY | 149 |
| All LYS Sites | ALY | 171 |
| All LYS Sites | ALY | 172 |
| All LYS Sites | ALY | 177 |
| All LYS Sites | ALY | 187 |
| All LYS Sites | ALY | 197 |
| All LYS Sites | ASN | 57 |
| All LYS Sites | ASN | 129 |
| All LYS Sites | ASP | 97 |
| All LYS Sites | HIS | 101 |
| All LYS Sites | ILE | 99 |
| All LYS Sites | ILE | 194 |
| All LYS Sites | LEU | 84 |

| | | |
|---|---|---|
| All LYS Sites | THR | 175 |
| All LYS Sites | VAL | 31 |
| All LYS Sites | PRO | 34 |
| All LYS Sites | PRO | 53 |
| All LYS Sites | PRO | 114 |
| All LYS Sites | PRO | 166 |

**Table 6. Residues with D-chirality involved in Condensate tau.**
